# Defining mucosal immunity using mass cytometry following experimental human pneumococcal challenge

**DOI:** 10.1101/546929

**Authors:** Simon P. Jochems, Karin de Ruiter, Carla Solórzano, Astrid Voskamp, Elena Mitsi, Elissavet Nikolaou, Beatriz F Carniel, Sherin Pojar, Esther L. German, Jesús Reiné, Alessandra Soares-Schanoski, Helen Hill, Rachel Robinson, Angela D. Hyder-Wright, Caroline M. Weight, Pascal F. Durrenberger, Robert S. Heyderman, Stephen B. Gordon, Hermelijn H. Smits, Britta C. Urban, Jamie Rylance, Andrea M. Collins, Mark D. Wilkie, Lepa Lazarova, Samuel C. Leong, Maria Yazdanbakhsh, Daniela M. Ferreira

**Affiliations:** Department of Clinical Sciences, Liverpool School of Tropical Medicine, Liverpool, United Kingdom; Department of Parasitology, Leiden University Medical Center, Leiden, Netherlands; Bacteriology Laboratory, Butantan Institute, Sao Paulo, Brazil; Royal Liverpool and Broadgreen University Hospital, Liverpool, United Kingdom; Division of Infection and Immunity, University College London, London, United Kingdom; Centre for Inflammation and Tissue Repair, University College London, London, United Kingdom; Malawi-Liverpool-Wellcome Trust Clinical Research Programme, Blantyre, Malawi; Department of Parasitology, Liverpool School of Tropical Medicine, Liverpool, United Kingdom; Aintree University Hospital NHS Foundation Trust, Liverpool, United Kingdom; Department of Otorhinolaryngology – Head and Neck Surgery, Aintree University Hospital NHS Foundation Trust, Liverpool, United Kingdom

**Keywords:** *Streptococcus pneumoniae*, Colonisation, Host-pathogen interaction, Mass cytometry, CyTOF, Controlled human infection, B cells, Mucosal immunology, mucosal associated invariant T cells, MAIT cells

## Abstract

*Streptococcus pneumoniae* (Spn) is a common cause of respiratory infection, but also frequently colonises the nasopharynx in the absence of disease. We used mass cytometry to study immune cells from nasal biopsy samples, collected following experimental human pneumococcal challenge, in order to identify immunological changes that follow and control spn colonisation. Using 37 markers, we characterized 293 nasal immune cell clusters, of which 7 were associated with Spn colonisation. B cell and CD8^+^CD161^+^ T cell clusters were significantly higher in non-colonised than in colonised subjects. Spn colonization led to recirculation of not only Spn-specific but also aspecific nasal B cells. This associated with increased numbers of circulating plasmablasts and increased antibody levels against the unrelated bacterium *Haemophilus influenzae*. In addition, we demonstrated that baseline functionality of blood mucosal associated invariant T (MAIT) cells associated with protection against Spn. These results identify new host-pathogen interactions at the mucosa upon Spn colonisation.

## Introduction

*Streptococcus pneumoniae* (Spn) is a major cause of morbidity and mortality worldwide (O’Brien et al., 2009; Welte et al., 2012). However, nasopharyngeal colonisation, or carriage, of Spn in the absence of disease is common, with approximately 50% of infants and 10% of adults colonised at any time (Goldblatt et al., 2005). Carriage is an immunising event in both children and adults but is also important as a prerequisite of disease and as the source of transmission (Ferreira et al., 2013; McCool et al., 2002; Melegaro et al., 2004; Simell et al., 2012).

Mouse models have suggested that Th17-mediated recruitment of neutrophils and monocytes to the nasopharynx is the mechanism of control and clearance of Spn carriage (Lu et al., 2008; Lu et al., 2010; Zhang et al., 2009). Recently, we demonstrated using an experimental human pneumococcal challenge (EHPC) model that carriage leads to degranulation of nasal-resident neutrophils and recruitment of monocytes to the nasal mucosal surface (Jochems et al., 2018). Protection against carriage acquisition was associated with the levels of circulating memory B cells, but not levels of IgG, directed against the Spn polysaccharide capsule (Pennington et al., 2016). However, the relative role of these and other adaptive and innate immune cell subsets in controlling Spn at the human nasal mucosa remains largely unknown (Khan and Pichichero, 2014).

Here, we applied mass cytometry (CyTOF) to nasal biopsies collected following experimental human pneumococcal challenge to comprehensively study mucosal immunity to Spn carriage.

## Results

### Characterization of nasal immune populations

Twenty healthy subjects negative for natural pneumococcal carriage at baseline screening were challenged intranasally with type 6B Spn (Fig. 1A and Table 1). Carriage state was assessed at days two and seven post challenge and a nasal biopsy was collected at ten days post challenge (Supplementary Video 1), the timepoint at which Spn starts to be cleared from the nose (Gritzfeld et al., 2014; Rylance et al., 2018). Eight subjects became colonized with Spn (carriage^+^), while twelve subjects remained carriage^−^ (Fig. 1A). Biopsies yielded a median of 2.3×10^5^ cells (IQR: 1.6×10^5^ - 3.2×10^5^) per subject, approximately 90% of which were stromal cells, which were stained with a panel of thirty-eight antibodies and analysed by CyTOF (Fig. 1B, Supplementary Table 1). Viable immune cells were manually gated from all acquired events and subsequently clustered by hierarchical-stochastic neighbour embedding (h-sne) using Cytosplore software (Fig. 1C,2) (Abdelmoula et al., 2018; Hollt et al., 2016; van Unen et al., 2017). H-sne is a recently developed method in which t-distributed stochastic neighbor embedding (t-sne) is performed sequentially to cluster first global cell populations, each of which is then in turn clustered into subpopulations.

**Table 1.** Volunteer

|  | Carriage<sup>-</sup><br>(n=12) | Carriage<sup>+</sup><br>(n=8) |
| --- | --- | --- |
| Female gender (%) | 4 (33.3%) | 4 (50%) |
| Median age (min-max) | 21 (18-44) | 23 (20-30) |
| Median day 2 Spn density CFU/mL (min-max) | - | 127.2 (5.8 - 38677.7) |
| Median day 7 Spn density CFU/mL (min-max) | - | 187.5 (0 – 21736.6) |

cohort characteristics divided by carriage state.

**Figure 1.**
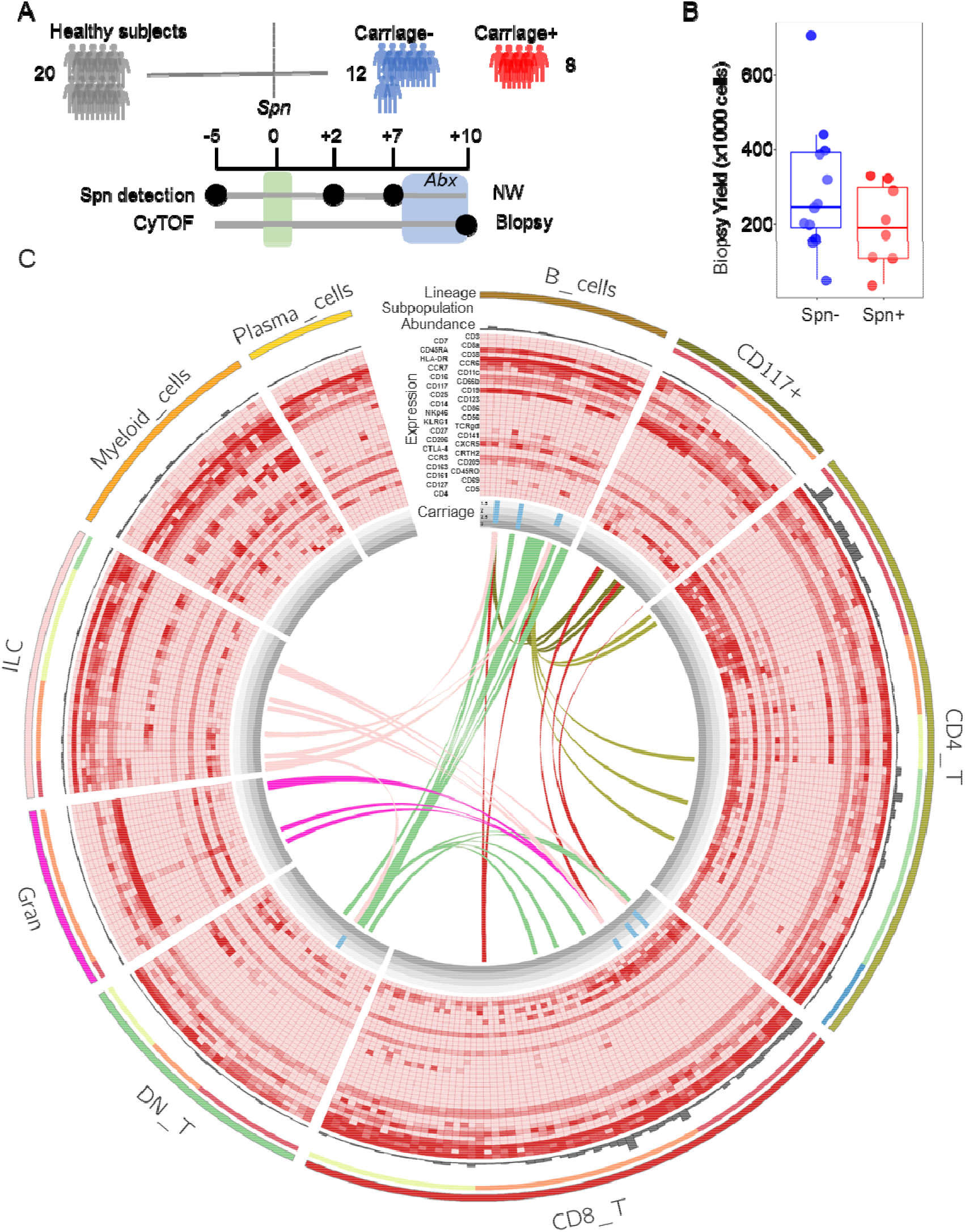
Mass cytometry from nasal biopsies following experimentabetween pneumococcal carriage and l human pneumococcal challenge. A) Study design showing pneumococcal inoculation (green bar) and sample collection. Subjects who acquired pneumococcus following challenge are depicted in red (n=8), while those protected are depicted in blue (n=12). Antibiotics (abx) were administered in the 3 days leading up to biopsy collection (blue area). B) Viable cell yield following enzymatic biopsy digestion for the twenty biopsies collected for CyTOF. Individual samples and boxplots are shown. C) Circle diagram showing all 293 defined clusters within 9 lineages and 22 subpopulations. From outside in: number of cells in each cluster is depicted by grey bars. Relative expression for 36 markers is shown with red depicting higher expression (CD45 and Epcam are not depicted). Association with carriage state is shown, where blue bars depict the fold-change of the median of normalized abundance in carriage^−^ subjects over carriage^+^ subjects. Ribbons connect highly correlated (r>0.70) clusters that were associated with Spn carriage status not belonging to the same lineage, with colour indicating the lineage of origin. DN_T = double negative T cells, Gran = granulocytes, ILC = innate lymphoid cells.

Based on the expression of 37 markers, a total of 199,426 immune cells from all subjects were divided into nine lineages (CD8^+^ T cells, CD4^+^ T cells, myeloid cells, innate lymphoid cells, B cells, double-negative T cells, granulocytes, CD117^+^ cells and plasma cells, in order of decreasing abundance). These cell lineages were further divided into twenty-two subpopulations and 293 clusters (Fig. 1C and Table 2). Cell numbers were normalized to the number of stromal cells for each subject to correct for varying biopsy yields. Normalized abundancies were then compared between carriage^−^ and carriage^+^ subjects for each of the lineages, subpopulations and clusters. There were no significant differences in frequencies between total lineages or subpopulations between carriage^−^ and carriage^+^ subjects. However, at a finer level seven clusters were significantly higher in carriage^−^ than in carriage^+^ subjects (Fig. 1C, blue bars). Of note, three B cell clusters were higher in carriage^−^ subjects (Fig. 1C). Moreover, three CD8^+^ T cell clusters, all expressing CD161, and one CD8^dim^ T cell cluster were higher in carriage^−^ subjects than in carriage^+^ subjects (Fig. 1C). The seven significant clusters strongly correlated (r>0.70) with eighty-eight clusters in other lineages/subpopulations, sixty-eight of which were in B or T cell lineages, highlighting an interconnectivity between B and T cell subpopulations in the human nasopharynx (Fig. 1C).

**Table 2.** List of lineages and subpopulations derived from nasal biopsy analysis. For all nine lineages and twenty-two subpopulations, the numbers of defined cell clusters are shown. In addition, the total numbers of cells within those lineages/subpopulations and the percentage of that subpopulation within all cells for carriage^−^ and carriage^+^ subjects are shown. Memory cells are defined as CD45RO^+^RA^−^ and naïve cells are defined as CD45RO^−^RA^+^

| Lineage | Subpopulation | Clusters | Cells | %Carriage<sup>-</sup> | %Carriage<sup>+</sup> |
| --- | --- | --- | --- | --- | --- |
| **CD8<sup>+</sup> T cells** | CD161<sup>+</sup> CD8<sup>+</sup> T cells | 17 | 25103 | 13.0 | 11.7 |
|  | Naïve CD8<sup>+</sup> T cells | 20 | 9042 | 3.8 | 6.3 |
|  | Memory CD8<sup>+</sup> T cells | 26 | 36860 | 18.1 | 19.4 |
|  | **Total** | **63** | **71005** | **34.9** | **37.3** |
| **CD4<sup>+</sup> T cells** | CD161<sup>+</sup> CD4<sup>+</sup> T cells | 21 | 36328 | 18.7 | 17.0 |
|  | CD25<sup>hi</sup> CD4<sup>+</sup> T cells | 9 | 2743 | 1.2 | 1.9 |
|  | Naïve CD4<sup>+</sup> T cells | 9 | 4235 | 2.2 | 1.8 |
|  | Memory CD4<sup>+</sup> T cells | 23 | 21158 | 10.5 | 10.8 |
|  | CD45RO<sup>-</sup>RA<sup>-</sup> CD4<sup>+</sup> T | 6 | 2743 | 1.2 | 1.9 |
|  | **Total** | **68** | **67207** | **33.8** | **33.5** |
| **Myeloid cells** | - | 25 | 15226 | 7.4 | 8.3 |
| **Innate lymphoid cells** | CD8<sup>-</sup>CD16<sup>+</sup> ILC | 13 | 3683 | 2.0 | 1.5 |
|  | CD8<sup>+</sup>CD16<sup>+</sup> ILC | 5 | 2392 | 1.1 | 1.4 |
|  | CD16<sup>-</sup>CD127<sup>+</sup> ILC | 4 | 1550 | 0.8 | 0.6 |
|  | CD16<sup>-</sup>CD127<sup>-</sup> ILC | 9 | 3417 | 1.7 | 1.8 |
|  | **Total** | **31** | **11042** | **5.6** | **5.3** |
| **B cells** | - | 22 | 10279 | 5.8 | 3.7 |
| **CD4<sup>-</sup>CD8<sup>-</sup> T cells** | TCRgd T cells | 9 | 1838 | 0.8 | 1.2 |
|  | DN T cells | 7 | 3052 | 1.6 | 1.5 |
|  | CD8<sup>dim</sup> T cells | 13 | 4317 | 2.2 | 2.1 |
|  | **Total** | **29** | **9207** | **4.6** | **4.7** |
| **Granulocytes** | CD66b<sup>+</sup> Granulocytes | 19 | 5478 | 2.9 | 2.4 |
|  | CD66b<sup>-</sup> Granulocytes | 2 | 662 | 0.3 | 0.3 |
|  | **Total** | **21** | **6140** | **3.2** | **2.7** |
| **CD117<sup>+</sup> cells** | CD117<sup>+</sup> lymphocytes | 8 | 2566 | 1.3 | 1.3 |
|  | CD117<sup>+</sup> mast cells | 13 | 2810 | 1.5 | 1.2 |
|  | **Total** | **21** | **5376** | **8.5** | **6.2** |
| **Plasma cells** | - | 13 | 3944 | 2.0 | 2.0 |
| **9** | **22** | **293** | **199426** | **100** | **100** |

### Nasal B cells are depleted during pneumococcal carriage

We then further investigated the three B cell clusters that were higher in carriage^−^ subjects (Fig. 3A,B). All three significantly higher clusters (cluster 4, 9 and 17) expressed CD45RA, HLA-DR, CD19, CCR6 and CCR7 to varying degrees. None of these clusters expressed CD38, a marker for plasmablasts, or CD5, a marker for innate B cells (Hardy, 2006; Jourdan et al., 2011). Cluster 9 was 2.9-fold higher in carriage^−^ subjects (p = 0.047) and cells in this cluster expressed also low levels of CXCR5 and CD27. Cluster 17 (2.0-fold higher, p = 0.049) additionally expressed the B cell activation marker CD69. To assess whether the higher frequency in carriage^−^ subjects was related to increased B cells in carriage^−^ subjects or decreased B cells in carriage^+^ subjects, we longitudinally measured CD19^+^ B cell frequencies in nasal microsamples collected from an independent cohort (Fig. 3C and Supplementary Fig. 1A). Compared to baseline, B cell levels decreased following pneumococcal carriage at days 2 (2.1-fold, p = 0.012), 6 (2.8-fold), 9 (2.0-fold) and 27 (3.1-fold, p = 0.007) post-inoculation. In the carriage^−^ group, B cell levels decreased 1.1-fold at days 2 and 6, increased 1.2-fold at day 9 and decreased 1.2-fold at day 27, respectively and were thus relatively stable.

**Figure 2.**
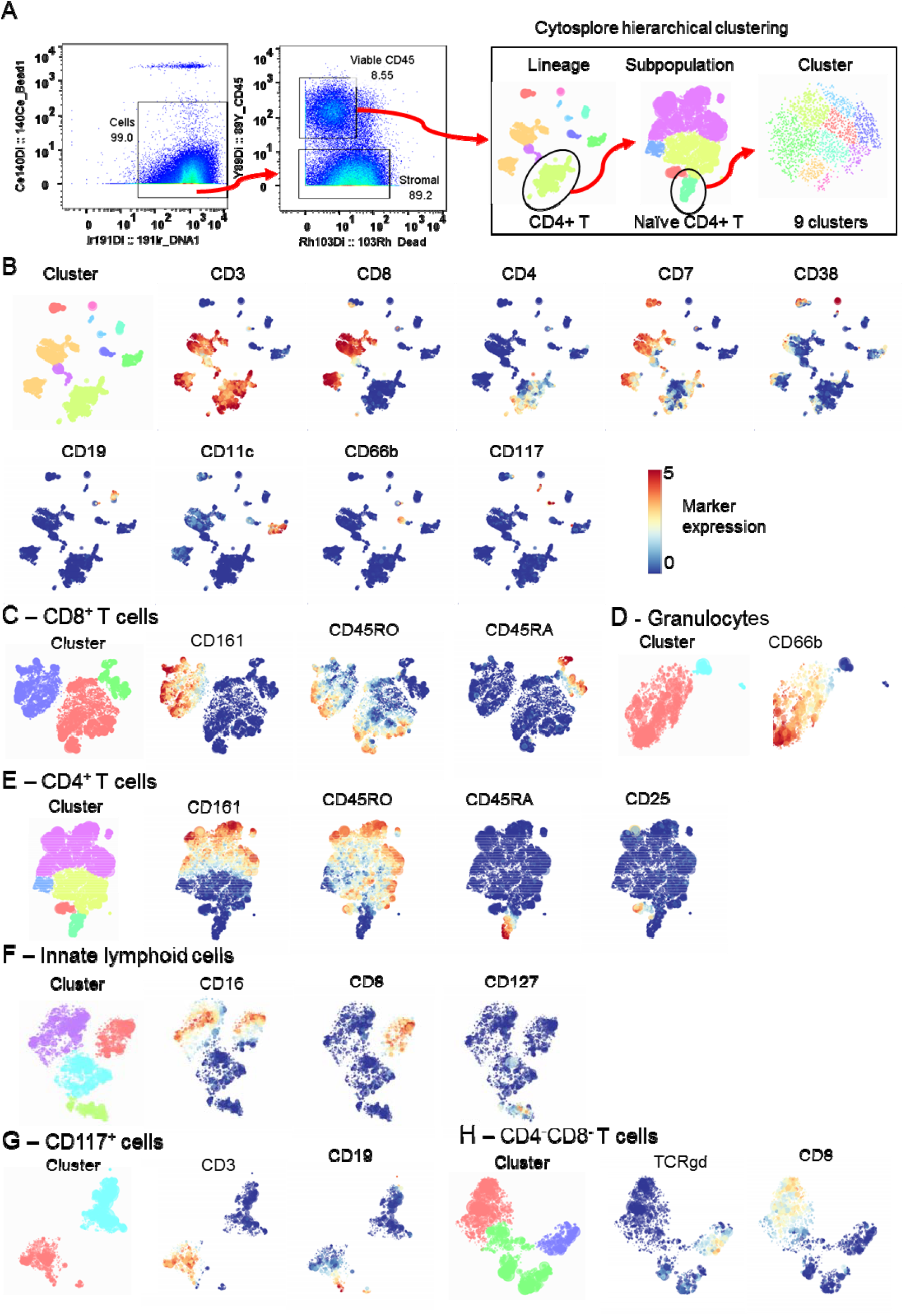
CyTOF analysis strategy. A) CyTOF data files were pre-gated using Flowjo to identify cells (DNA^+^ Bead^−^), followed by selecting viable immune cells (CD45^+^ Dead^−^). These cells were exported and loaded in Cytosplore for hierarchical stochastic neighbour embedding (h-sne), in which lineages, subpopulations and clusters were sequentially identified in three steps. Gating for naïve CD4^+^ T cells is shown as an example. B) Cells were clustered using all 38 markers minus the epithelial marker Epcam and lineages were then defined based on the expression of nine markers. Clustered lineages and expression of included markers are shown. Subpopulations for C) CD8^+^ T cells, D) granulocytes, E) CD4^+^ T cells, F) innate lymphoid cells, G) CD117^+^ cells and H) double-negative T cells were defined based on the expression of the depicted markers. B cells, plasma cells, myeloid cells were not further divided into subpopulations due to lack of clear clustering by relevant markers. Cell subpopulations were then further divided into clusters using all 38 markers minus the epithelial marker Epcam.

**Figure 3.**
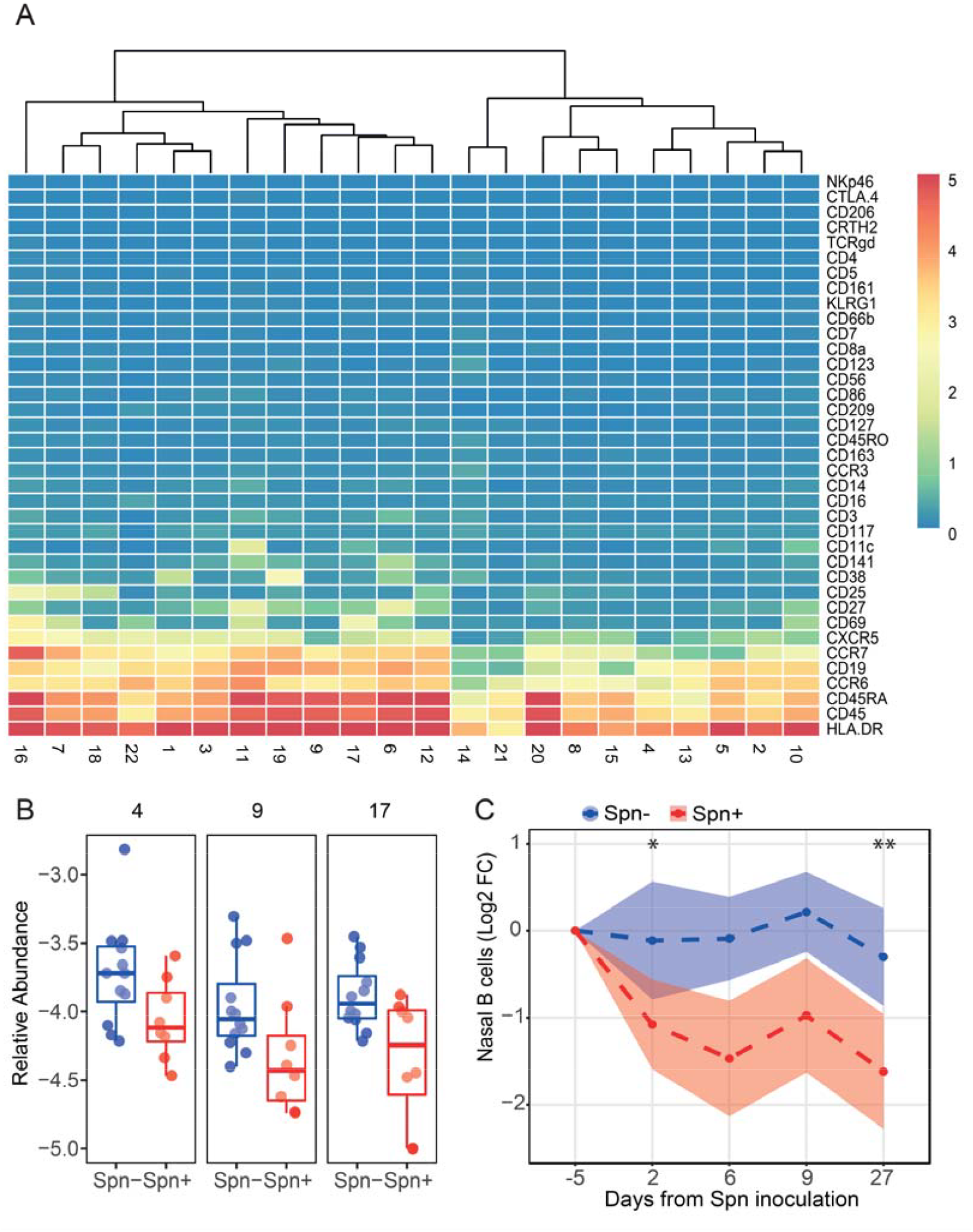
Nasal B cells are depleted following pneumococcal carriage. A) Heatmap showing the expression of thirty-seven markers for all B cell clusters. Clusters were ordered based on similarity and a distance dendrogram is depicted. B) The relative abundance for each of the three significantly higher clusters normalized to stromal cells is expressed on a log_10_ scale for carriage^−^ (Spn–, blue) and carriage^+^ (Spn+, red) subjects. Boxplot and individual subjects are depicted. C) Levels of nasal B cells longitudinally measured by flow cytometry from minimally-invasive nasal curettes in an independent cohort for carriage^−^ (Spn–, blue, n=52) and carriage^+^ (Spn+, red, n= 42) subjects. Mean and standard error of mean of log2-transformed fold change levels to baseline are shown. * p < 0.05, ** p < 0.01 by Wilcoxon test comparing to baseline.

### Pneumococcal carriage increases circulating plasmablasts

We hypothesized that the depletion of B cells from the nasal mucosa following carriage establishment was due to a re-circulation of activated B cells. Numbers of Spn-specific and total plasmablasts were measured in peripheral blood mononuclear cells (PBMC) collected before and after carriage establishment using a flow cytometry-based assay (Supplementary Fig. 1B). During carriage, the frequency of 6B polysaccharide-specific plasmablasts among total B cells increased while the frequency of plasmablasts specific to the pneumococcal protein pneumolysin remained unaltered (Fig. 4A). As a negative control we measured levels of plasmablasts specific for an unrelated Spn capsular type (15B), which were not affected as expected. However, the frequency of total circulating plasmablasts among all B cells increased (median 1.5x, IQR: 1.2-2.8x; p = 0.008) suggesting that nasal B cells became non-specifically activated during carriage. Similar results were obtained when normalizing to the total number of lymphocytes, demonstrating this was not due to other shifting B cell populations (Supplementary Fig. 2A). We then investigated CCR10 expression on these plasmablasts, which has been reported to mark IgA secreting cells (Morteau et al., 2008) and is potentially important for homing of B cells to mucosal tissues including the airways (Kato et al., 2013; van Splunter et al., 2018). The total population of plasmablasts post carriage displayed reduced numbers of CCR10^+^ cells, in contrast to 6B-specific plasmablasts, indicating differential expansion between specific and non-specific B cell populations (Fig. 4B). This is supported by the observation that increased circulating levels of 6B polysaccharide-specific plasmablasts inversely correlated with the nasal B cell CyTOF clusters 9 and 20, while total plasmablast increases inversely correlated with the CyTOF B cell clusters 21 (Fig. 4C). Clusters 9 and 21 still negatively correlated with levels of circulating 6B-specific and total plasmablasts, respectively, after normalization to total lymphocyte numbers (Supplementary Fig. 2B). Thus, it is likely that both polysaccharide-specific as well as unrelated B cells became activated following carriage, leading to recirculation. We then measured antibody levels in serum against not only Spn but also *Streptococcus pyogenes, Staphylococcus aureus* and *Haemophilus influenzae* as these are common colonizers of the human nasopharynx and thus nasal B cells against these bacterial species are likely present in the nose of most individuals. Following Spn colonisation, IgG levels specific for Spn (median 1.4x, IQR: 1.1-2.4) and *Haemophilus influenzae* (median 1.2x, IQR: 1.1-1.5) significantly increased, while IgG levels specific for *Streptococcus pyogenes* and *Staphylococcus aureus* were not significantly altered (Fig. 4D). Consistent with the decrease in CCR10 levels on total plasmablasts but not on 6B-specific plasmablasts, serum levels of specific IgA only increased for Spn and not for *Haemophilus influenzae* or any of the other bacterial species (Fig. 4E).

**Figure 4.**
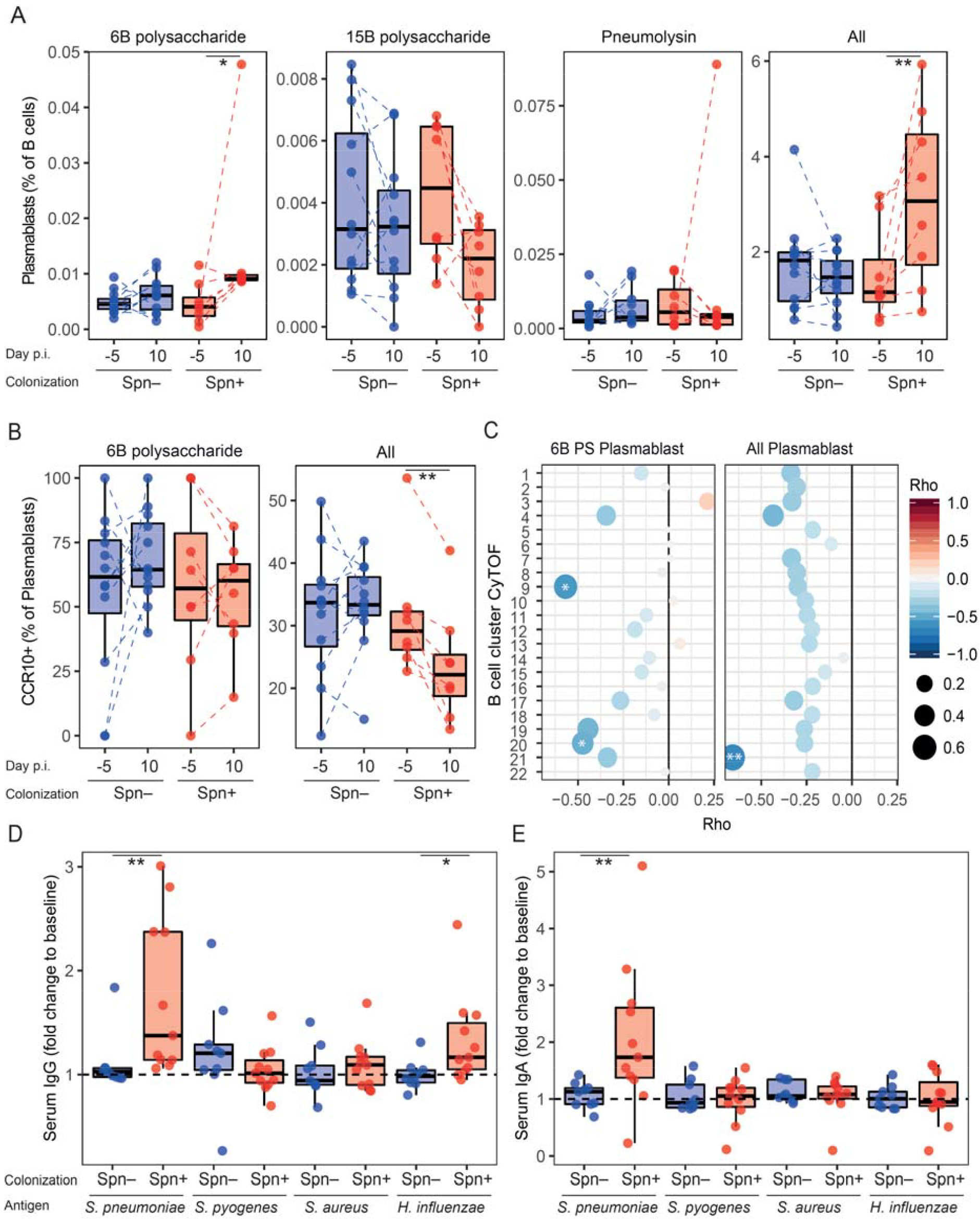
Pneumococcal carriage leads to increased systemic plasmablasts. A) Levels of 6B polysaccharide-specific, 15B polysaccharide-specific, Pneumolysin derivative b (Pneumolysin)-specific or all plasmablast amongst total B cells were measured from PBMC collected at baseline (Day -5) and at the time of biopsy (Day 10 post inoculation). Boxplots and individual subjects are depicted with carriage^−^ in blue (n=12) and carriage^+^ in red (n=8), with paired samples connected by dashed lines. * p < 0.05, ** p < 0.01 by Wilcoxon test comparing a group to its baseline. B) Levels of CCR10^+^ plasmablasts for 6B-specific and total plasmablasts measured from PBMC collected at baseline (Day -5) and at the time of biopsy (Day 10 post inoculation). Boxplots and individual subjects are depicted with carriage^−^ in blue and carriage^+^ in red with paired samples connected by dashed lines. ** p < 0.01 by Wilcoxon test comparing a group to its baseline. C) Correlations between fold-change in levels of 6B PS-specific and total plasmablasts between baseline and day 10 against levels of B cell clusters measured by CyTOF. Color and size of symbols reflect the Spearman rho value. * p < 0.05 and ** p < 0.01 by Spearman test. Fold change (day 23 post inoculation versus baseline) in levels of D) IgG and E) IgA against *Streptococcus pneumoniae, Streptococcus pyogenes, Staphylococcus aureus* and *Haemophilus influenzae*. Boxplots and individual subjects are depicted with carriage^−^ in blue (Spn–, n=9) and carriage+ in red (Spn+, n=11). * p < 0.05, ** p < 0.01 by Mann-Whitney test comparing fold-change levels between carriage^−^ and carriage^+^ subjects.

### Nasal CD8 Tissue-resident memory T cells are higher in carriage^−^ subjects

The three clusters of CD8^+^ T cells and the cluster of CD8^dim^ T cells that were higher in carriage^−^ subjects all expressed CD69, a marker of tissue-resident memory (Trm) cells (Fig. 5A). To verify that these CD69^+^ CD8^+^ T cells represented Trm cells, we measured the expression of CD103 and CD49a on CD69^+^ and CD69-cells by flow cytometry from a representative biopsy (Supplementary Fig. 3A). Indeed, 89.1% of nasal CD69^+^ CD8^+^ T cells expressed CD103 and CD49a, confirming that these were Trm cells (Fig. 5B) (Kumar et al., 2017). The markers CD5, CD38, HLA-DR, CCR6, CD127, CCR7 and CD11c were expressed in cluster-specific patterns and at varying intensities among the significant clusters. This suggests that clusters of cells with varying degrees of activation and memory types were enriched in carriage^−^ subjects. One cluster expressed only low levels of CD8 (cluster 10 of CD8^dim^ T cells, 2.0-fold higher, p = 0.016), which could reflect cytotoxic effector memory cells (Trautmann et al., 2003). We then stimulated nasal biopsy cells and PBMC overnight with PMA and ionomycin to assess the functional capacity of nasal CD8^+^ T cells (Fig. 5C). Among nasal CD8^+^ T cells, 94.8% produced tumor necrosis factor alpha (TNF) and/or interferon gamma (IFN-□) following stimulation, compared to 36% of blood CD8^+^ T cells, demonstrating that nasal CD8^+^ T cells are highly functional.

**Figure 5.**
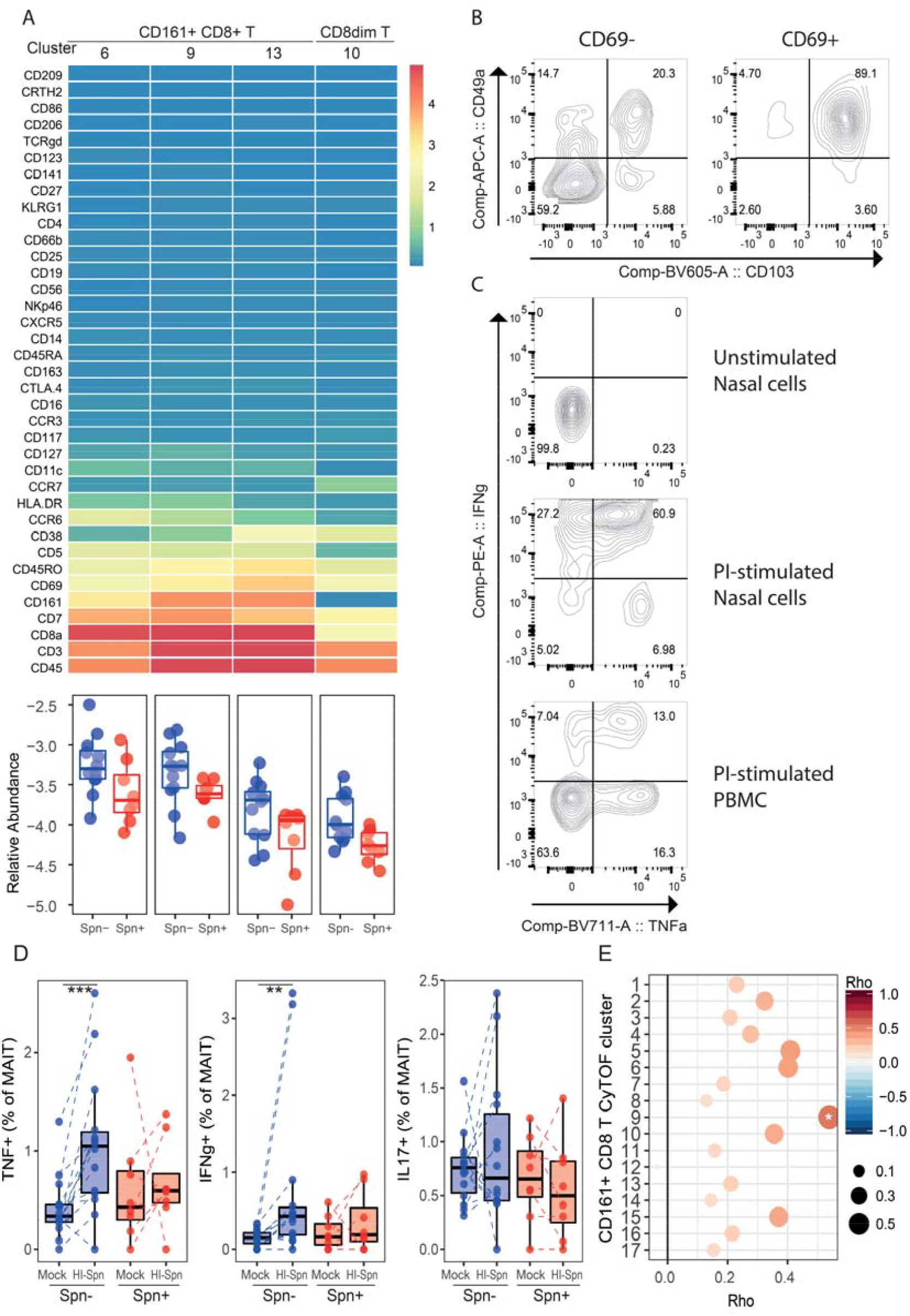
Increased MAIT responses associate with protection from carriage. A) Heatmap showing the expression of thirty-seven markers for each of the four CD8^+^ clusters that were significantly different between carriers and non-carriers. Non-significant CD8^+^ T clusters are not shown. Below the heatmap, the abundance for each of the significantly higher clusters normalized to stromal cells is expressed on a log_10_ scale for carriage^−^ (blue) and carriage^+^ (red) subjects. Boxplot and individual subjects are depicted. B) Representative flow cytometry contour plot of CD8^+^ CD69^+^ and CD8^+^ CD69^-^ T cells, showing CD103 and CD49a tissue resident marker expression on nasal biopsy cells. C) Representative flow cytometry contour plot of unstimulated nasal biopsy cells, and nasal biopsy cells and PBMC stimulated overnight with PMA and ionomycin (PI) to assess functional capacity. D) TNF, IFN-□and IL-17A production by CD8^+^ MAIT cells (CD161^+^TCRvα7.2^+^) after 3-day *in vitro* stimulation with heat-inactivated pneumococcus (HI-Spn) or left unstimulated for carriage^−^ (blue, n=14) and carriage+ (red, n=8) subjects in PBMC collected at baseline. Boxplots and individual subjects, connected by dashed lines, are shown. ** p < 0.01 by Wilcoxon test, *** p < 0.001 by Wilcoxon test. E) Correlations between the difference in cytokine production (total of TNF and IFN-□) by MAIT cells in vitro stimulated with HI-Spn or left unstimulated against CD8^+^ CD161^+^ T cell clusters measured by CyTOF. Colour and size of symbols reflect the Spearman rho value. * p < 0.05 by Spearman test.

### Baseline circulating MAIT functionality associates with resistance to pneumococcal carriage

Three of the four significant clusters expressed CD161, a marker for mucosal associated invariant T (MAIT) cells, and we therefore tested the hypothesis that MAIT cell responses against Spn were associated with protection against carriage. PBMC collected prior to pneumococcal challenge were stimulated *in vitro* with heat-inactivated Spn and activation (CD69) and cytokine production (TNF, IFN-□and IL-17A) were assessed (Supplementary Fig. 3B). MAIT cells of both carriage^−^ and carriage^+^ groups upregulated CD69 after a 3-day culture with heat-inactivated Spn (Supplementary Fig. 3C). However, only MAIT cells from carriage^−^ subjects produced increased levels of TNF and IFN-□, but not IL-17A, upon restimulation in vitro with heat-inactivated Spn (Fig. 5D). Conversely, MAIT cells from carriage^+^ subjects did not produce increased levels of any cytokine upon stimulation. This was specific to MAIT cells as conventional CD8^+^ T cells responded by producing small amounts of IFN-□and no TNF (Supplementary Fig. 3D). The baseline responses of MAIT cells in blood upon restimulation showed a positive correlation with numbers of nasal cells at ten days post pneumococcal challenge in CyTOF CD161^+^ CD8^+^ T cell cluster 9, which was significantly higher in the carriage^−^ group (r = 0.54, p = 0.02, Fig. 5E).

### Nasal monocytes show limited differentiation into macrophages

Monocytes have been previously associated with the clearance of Spn carriage (Jochems et al., 2018; Zhang et al., 2009), however these cells have not been previously phenotyped in detail in the human nasopharynx. Of the twenty-five clusters defined in the myeloid lineage, fifteen expressed CD14 (Supplementary Fig. 4). Of these, only two also expressed CD16. Four CD14^+^ clusters expressed the macrophage markers CD163 and CD206 and an additional three clusters expressed CD206 but not CD163 (Kaku et al., 2014). However, alveolar monocytes can express CD206, suggesting this is not a definitive indication of differentiation (Yu et al., 2016). The activation markers CD25 and CD86 were present on five monocyte clusters (Farina et al., 2004). Thus, monocyte/macrophages in the nose mainly consisted of classical monocytes with limited differentiation into macrophages.

### Characterization of nasal CD4^+^ memory T cells

CD4^+^ T memory cells, in particular Th17 cells, were previously found to be critical for Spn immunity in mice models of nasal colonisation (Lu et al., 2008; Zhang et al., 2009). Of all cells in the CD4^+^ T cell lineage, 89.6% expressed the memory marker CD45RO. Of these, 60.3% expressed CD161, a marker that has been proposed to identify Th17 cells (Cosmi et al., 2008; Kleinschek et al., 2009). Another 4.6% of memory cells was defined by expression of high levels of CD25, a marker for regulatory T cells. We defined twenty-three clusters of CD161^−^ CD4^+^ T memory cells, twenty-one clusters of CD161^+^ CD4^+^ T memory cells and nine clusters of CD25^hi^ CD4^+^ T memory cells (Supplementary Fig. 5). All CD4^+^ T memory cell clusters expressed a combination of the markers CD7, CD127, HLA-DR, CD38 and CD69 demonstrating a wide range in activation and differentiation status (Kumar et al., 2017). The CD25^hi^ CD4^+^ T memory cells likely were regulatory T cells as they were predominantly negative for CD127 and two of these clusters expressed cytotoxic T-lymphocyte-associated protein 4 (CTLA-4) and CD27 (Ballke et al., 2016). CD161 was not restricted to Th17 cells as among CD161^+^ CD4^+^ T memory cells, two clusters expressed also CD8 and were thus double-positive T cells (Overgaard et al., 2015). In addition, two clusters expressed CD25 without CD127 expression indicating regulatory T cells, and one cluster expressed chemoattractant receptor-homologous molecule expressed on Th2 cells (CRTH2), a marker of Th2 cells (Cosmi et al., 2000).

### Cellular distribution through the nasal mucosa

We then performed immunohistochemistry on a biopsy from a challenged but carriage^−^ subject to further understand the distribution of these cells through the mucosal tissue (Fig. 6). CD4^+^ T cells were found predominantly in the subepithelial layer (Fig. 6C,D), while CD8 and CD161 were also found at the epithelial layer (Fig. 6E,F). Similar to CD4^+^ T cells, B cells (defined by CD20) were mostly observed in the sub-epithelium, while myeloid cells (CD68) could be seen at both the epithelial and sub-epithelial layer (Fig. 6G,H). Neutrophils were found abundantly at the epithelial surface but also in the sub-epithelium (Fig. 6I,J).

**Figure 6.**
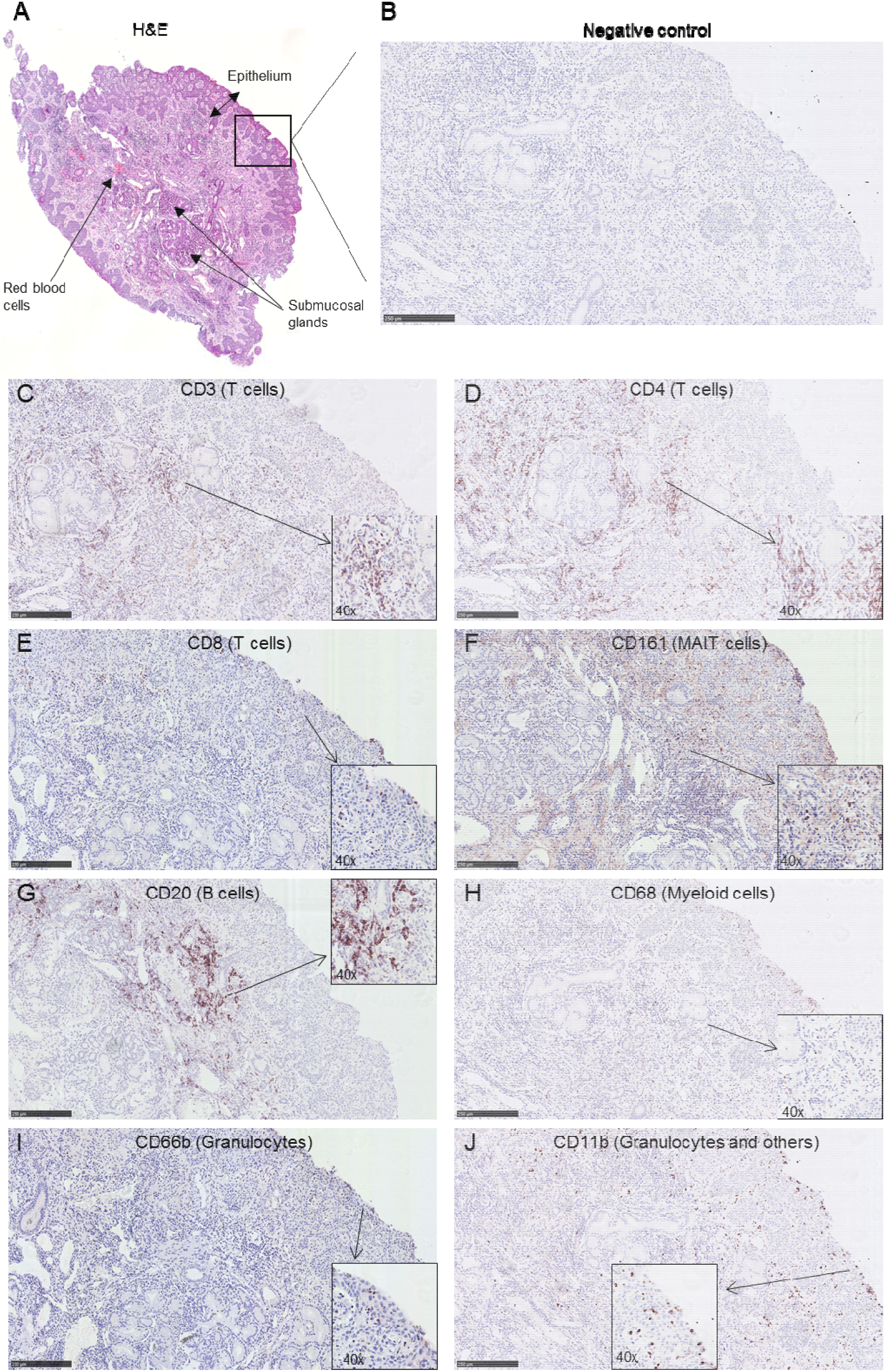
Immunohistochemistry on serial sections of a nasal biopsy. To establish an overall cellular distribution in the tissue a 10x magnification is shown for each of the markers. A 40x inset is also included to visualize some individual positive cells. A) Haematoxylin and eosin staining showing the entire biopsy. Staining of subsequent slices showing the biopsy at the epithelial edge for the markers B) negative control, C) CD3, D) CD4, E) CD8, F) CD161, G) CD20, H) CD68, I) CD66b and J) CD11b. A scale showing 250µm are added to all panels and a 40x inset is included. Slices were counterstained with haematoxylin and eosin. Some background staining of the extracellular matrix is present for CD161 (panel F).

## Discussion

This study comprehensively characterised immune cells in biopsies collected from the human nasal mucosa. As nasal samples were collected ten days following experimental human pneumococcal challenge, we were able to associate the frequency of specific immune populations with Spn carriage. Given the difficulty in access to such tissue samples, especially in a setting where the onset of infection is known, this provided a unique opportunity to investigate mucosal immune responses not undertaken previously. The application of CyTOF led to a broad and comprehensive study of cellular subsets involved in immunity against Spn carriage, deriving 293 immune clusters belonging to nine cellular lineages. Clusters belonging to B cells and CD8^+^ CD161^+^ T cells were higher in carriage^−^ subjects.

B cells were depleted from the nasal mucosa following the establishment of Spn carriage. This depletion correlated on an individual level with increased numbers of circulating 6B polysaccharide specific and total plasmablasts. Thus, this depletion likely was due to recirculation of activated B cells rather than due to apoptosis of nasal B cell upon Spn polysaccharide capsule encounter as has also been described (Dullforce et al., 1998). The total plasmablast expansion, but not 6B plasmablast expansion, was characterized by a decreased proportion of CCR10^+^ cells, suggesting a preferential expansion of CCR10^-^ cells or a downregulation of this marker. The correlation between low numbers of cells in specific nasal B cell clusters with increased levels of circulating plasmablasts indicates that activation of nasal B cells during carriage led to B cell re-circulation. Indeed, trafficking of memory B cells between airways and blood has been reported (Ohm-Laursen et al., 2018).

As B cells express the innate receptors TLR2 and TLR4 (Hayashi et al., 2005), which can be activated by pneumococcus, it is possible that carriage leads to non-specific B cell activation. This in turn could lead to increased protection against unrelated pathogens following pneumococcal carriage by boosting humoral responses. Indeed, levels of serum IgG against *Haemophilus influenzae* increased following colonization with Spn. We observed no increase in serum IgG levels against *Streptococcus pyogenes* or *Staphylococcus aureus*. It is possible that a limited number of pre-existing nasal B cells against these bacteria were present in the nose of these individuals at the time of colonization as their previous colonization history, and longevity of mucosal B cells, are unknown. It is also not impossible that the whole cell ELISA used here was unable to detect an increase against specific bacterial antigens. Negative associations between colonisation with Spn and *Haemophilus influenza* have been frequently reported and these are thought to rely on direct bactericidal effects of bacteria, competition for host nutrients and adherence factors, or activation of innate immune responses (de Steenhuijsen Piters et al., 2015). Our results suggest that non-specific activation of B cells could be another mechanism by which species within the microbiota interact with each other indirectly. *Neisseria lactamica* has been previously demonstrated to be able to aspecifically activate innate B cells, suggesting this is not a unique feature of Spn (Vaughan et al., 2010; Vaughan et al., 2009)

Several nasal CD8^+^ Trm cell clusters were higher in subjects protected from Spn carriage. These cells were previously found to be protective against influenza infection in murine models (Pizzolla et al., 2017). Spn is classically thought of as an extracellular bacterium and therefore the role of CD8^+^ T cells in controlling Spn has not been extensively studied in humans. However, it was recently shown that Spn can replicate within splenic macrophages and can reside within epithelial cells, suggesting that CD8^+^ T cell immunity could be elicited by Spn and play a role in protection against Spn carriage or disease (Ercoli et al., 2018; Weight et al., 2018). Indeed, Spn protein specific CD8^+^ T cells could be readily detected in blood of Gambian adults (Mureithi et al., 2009). In murine models, depletion of CD8^+^ T cells was protective against Spn lung infection but did not have an effect on nasopharyngeal carriage (Malley et al., 2005; Weber et al., 2011).

We found here that CD8^+^ MAIT cell functionality before pneumococcal challenge associated with a resistance to carriage acquisition. MAIT cells are a recently identified T cell subset that is common in humans, consisting of up to 10% of all T cells in the circulation, but that is very rare in mice (Wakao et al., 2017). It is possible that this difference has led to an underappreciation of the CD8+ T cell’s role in protection against pneumococcal carriage in humans. MAIT cells were recently reported to be able to recognize Spn through MHC class I-related protein 1 (MR-1) dependent and independent pathways (Kurioka et al., 2018). MAIT cells were previously found to be important in the protection against lung bacterial and viral infections via direct and indirect responses (Wakao et al., 2017). Our findings now suggest these cells could also protect against nasopharyngeal Spn colonisation. Baseline MAIT functionality in blood positively correlated with cell numbers within one of the nasal CD8^+^ CD161^+^ cell clusters, suggesting trafficking of MAIT cells from the blood to the nose upon pneumococcal encounter. Indeed, MAIT cells have been shown to be depleted from the circulation and accumulate in tissues upon infection (Chen et al., 2017; Howson et al., 2018).

One limitation of this study is that the number of granulocytes measured was very low due to the overnight resting step following enzymatic digestion. While this resting step allowed for the return of markers that were cleaved by the enzymatic digestion, neutrophils quickly become apoptotic after being removed from the body (Autengruber et al., 2012; Goodyear et al., 2014; Pongracz et al., 1999). Consequently, the characterization of granulocytes reported here is incomplete and we were not able to assess whether specific neutrophil subsets are associated with protection against pneumococcal colonisation. In addition, due to the invasiveness of sample acquisition, sample size was limited and we were not able to characterize nasal biopsies at various time points. Thus, no baseline was available and transient responses early after bacterial inoculation could not be assessed. As subjects received antibiotics prior to biopsy collection, we were unable to associate levels of any of the monocyte or CD4^+^ T cell clusters with Spn clearance.

In conclusion, this study provides both a broad and an in-depth view of the adult human nasal immune system in the setting of experimental human pneumococcal challenge. Nasal B cells were depleted following carriage establishment, likely due to recirculation following non-specific and specific activation. This associated with increases in antibody responses against not only Spn, but also *Haemophilus influenzae*. In addition, CD8^+^ MAIT cell responses were associated with protection from Spn carriage.

## Acknowledgements

This work was supported by the Medical Research Council (grant MR/M011569/1) to SG, and by support from Bill and Melinda Gates Foundation (grant OPP1117728) and the National Institute for Health Research (NIHR) Local Comprehensive Research Network to DF. This work was supported by the Human Infection Challenge Network for Vaccine Development (HIC-Vac) funded by the GCRF Networks in Vaccines Research and Development which was co-funded by the MRC and BBSRC. RSH and CW are funded through the NIHR Global Health Research Unit on Mucosal Pathogens at UCL. Flow cytometric acquisition was performed on a BD LSR II funded by a Wellcome Trust Multi-User Equipment Grant (104936/Z/14/Z). Purified pneumococcal Pneumolysin derivative b (Pdb) protein was a kind gift by Dr. Eliane Miyaji. We would like to thank all volunteers for participating in this study and C. Lowe, C. Hales, H. Adler, V. Connor, C.J. Webb and A. Panarese for clinical support.

## Author contributions

SJ contributed to conceiving, designing, conducting and analysing experiments, design of the study and writing of the paper. KR, CS, AV contributed to designing, conducting and analysing experiments. SG, LL, JRylance, AC, SL contributed to conceiving and designing the study. EM, EN, BC, AS, SP, EG, JReine, CW and PD contributed to conducting and analysing experiments. HH, RR, AHW, SL and MW contributed to sample collection. RSH, HS, BU and MY contributed to designing and analysing experiments. DF contributed to conceiving, designing and analysing experiments, design of the study and writing of the paper. All authors have read and approved the manuscript.

## STAR Methods

### Study design and sample collection

Healthy adult subjects were screened for the presence of natural pneumococcal carriage in nasal wash samples (NW) using classical microbiology (Ferreira et al., 2013; Gritzfeld et al., 2014; Gritzfeld et al., 2013). Subjects not naturally carrying pneumococcus were then inoculated with 80,000 CFU per nostril of 6B type Spn as described (Ferreira et al., 2013; Gritzfeld et al., 2013). Development of nasal carriage was monitored using NW samples collected at days 2 and 7 post inoculation. Growth of pneumococcus from NW samples at any time-point defined carriage positive volunteers. All subjects then received a three-day course of amoxicillin and underwent a 4mm nasal biopsy at day 10 post inoculation. The nasal cavity was first sprayed up to six times with lidocaine hydrochloride 5% with phenylephrine hydrochloride 0.5%. Five to ten minutes later the infero-medial part of the inferior turbinate, i.e. the point of incision, was injected with up to 1 mL of lidocaine hydrochloride 2% with adrenaline 1:80 000. An incision of approximately 5 mm with No.15 blade was then made and 2-4 mm of mucosal tissue was removed with Tillies Henckle’s surgical forceps. This study was registered under ISRCTN85509051. Nasal curettes (ASL Rhino-Pro©, Arlington Scientific) were collected from a second cohort (ISRCTN16993271) of inoculated subjects as previously described. The outcomes reported in this manuscript were a priori included in the study protocols.

### Ethics statement

All subjects gave written informed consent and research was conducted in compliance with all relevant ethical regulations. Ethical approval was given by the East Liverpool NHS Research and Ethics Committee (REC), reference numbers: 17/NW/0029 and 14/NW/1460.

### Nasal biopsy digestion

Nasal biopsies were finely cut using a sterile scalpel size 11 (Fisher Scientific). Pieces were then incubated in 20mL pre-warmed RPMI 1640 (Fisher Scientific) with Liberase TL (250μg/mL, Sigma) and DNAse I (50μg/mL, Sigma). Fragments were incubated for 45 minutes at 37°C, while shaking at 250rpm at a 10° angle. At the end of the digestion, biopsies were passed five times through a 16-gauge blunt-ended needle (Fisher Scientific) and the digested sample was filtered over a 70um filter (Fisher Scientific). This process was repeated for any remaining fragments. Cell were spun down for 10 minutes at 400xg and then red blood cells were lysed using an osmotic lysis buffer. Cells were washed with RPMI with 20% heat-inactivated fetal bovine serum (FBS, Fisher Scientific), resuspended at 10^6^ cells/mL in RPMI with 20% FBS and rested overnight. The next day, cells were counted and washed with RPMI + 10% FBS. Cells were stained as a viability marker using 1µM intercalator Rh-103 (Fluidigm) for 15 minutes, washed and fixed with 1.8% paraformaldehyde (Sigma) for 15 minutes. Cells were washed and stored in liquid nitrogen in CTL-Cryo™ ABC media (Cellular Technology Limited) until CyTOF barcoding and staining.

### Mass cytometry staining and analysis

Nasal biopsy cells were thawed on ice and barcoded using the Cell-ID 20-plex Pd Barcoding Kit as per manufacturer’s instructions (Fluidigm). The effect of fixation on epitopes detected by the included antibody clones was tested using PBMCs and monocyte-derived dendritic cells. Following three washes with staining buffer (Fluidigm) and 10 minutes of FcR blocking (Biolegend) pooled cells were stained for 45 minutes at room temperature with the antibody cocktail (Supplementary Table 1). Cells were washed twice with staining buffer and incubated for 1 hour with 1000x diluted 125 μM Cell-ID intercalator-Ir (Fluidigm) to stain DNA. Cells were washed 3 times with staining buffer and 2 times with de-ionized H_2_O prior to addition of normalization beads (Fluidigm) and acquisition on a Helios 2 mass cytometer (DVS Sciences). CyTOF Fcs files were normalized using the included beads, concatenated and debarcoded as per manufacturer’s instructions. The debarcoding step leads to a removal of doublets (Zunder et al., 2015). Then, viable immune cells were pre-grated (Fig. 2) and exported as .fcs files using Flowjo X (Treestar). These were further analysed using Cytosplore (https://www.cytosplore.org/).

### Nasal B cell phenotyping

Immunophenotyping of nasal B cells obtained by curettes was performed as described (Jochems et al., 2017). In brief, cells were dislodged from curettes and stained with LIVE/DEAD® Fixable Aqua Dead Cell Stain (ThermoFisher) and an antibody cocktail containing among others Epcam-PE, HLADR-PECy7, CD66b-FITC, CD19-BV650 (all Biolegend), CD3-APCCy7, CD14-PercpCy5.5 (BD Biosciences) and CD45-PACOrange (ThermoFisher). Samples were acquired on a LSRII flow cytometer and analysed using Flowjo X (Treestar). Fluorescent minus one controls for each of the included antibodies were used to validate results during set-up of all of the panels used. Samples with less than 500 immune cells or 250 epithelial cells (13.3% of all nasal samples) were excluded from further analysis.

### Intracellular cytokine staining following PMA/Ionomycin or pneumococcus stimulation

For intracellular cytokine staining after PMA and Ionomycin stimulation, fresh nasal biopsy cells or PBMC were stimulated with 100 and 500 ng/mL of these, respectively. After 2 hours, Golgiplug™ (BD Biosciences) was added and cells were incubated for another 16 hours. Cells were washed and stained extracellularly with LIVE/DEAD® Fixable Violet Dead Cell Stain (ThermoFisher) for 15 minutes and then for another 15 minutes with CD161-APC, CD69-BV650, CD25-PEDazzle594, CD103-BV605, CD4-PercpCy5.5, CD8-AF700, TCRvα7.2-BV785 (all Biolegend) and CD3-APH7 and TCRgd-PECy7 (BD Biosciences). Cells were then permeabilized using the eBioscience™ Foxp3 / Transcription Factor Staining Buffer Set (Fisher Scienctific) following the manufacturer’s protocol. Intracellular staining was done for 30 minutes with FOXP3-AF488, IFNg-PE, TNFa-BV711 (Biolegend) and IL17A-BV510 (BD Biosciences). Finally, cells were washed, resuspended in 200µL PBS and acquired on a LSR2.

For staining with pneumococcus, PBMC were thawed with 50μg/mL DNAse I (Sigma) in pre-warmed RPMI + 10% FBS and washed twice, once in media including DNAse I and once in media without DNAse I. Cells were rested overnight and then cultured at 5×10^5^ cells in 500uL media with 5ug/mL (corresponding to 4.3×10^7 CFU/mL) heat-inactivated type 6B *Streptococcus pneumoniae* or left unstimulated as a control. After 48 hours, fresh antigen was added to the cells and 2 hours later Golgiplug was added and cells were treated as above.

### Pneumococcal-specific B cell detection

Purified pneumococcal polysaccharides 6B and 15B (Oxford Biosystems) and Pdb were diluted to 100µg/mL in purified H_2_O and biotinylated using the One-Step Antibody Biotinylation Kit (Miltenyi) as per manufacturer’s instructions. Biotinylated proteins were then 2x dialysed for 45 minutes against 1L PBS using Slide-A-Lyzer™ MINI Dialysis Device, 3500 molecular weight cut off (ThermoFisher) and stored at 4°C until labelling. Biotinylated 15B, 6B and Pdb were then mixed in a 4:1 molecular ratio (Pdb), or a 1:1 molecular ratio (polysaccharides), with PE-streptavidin, BV785-streptavidin or FITC-streptavidin (Biolegend), respectively. Incubation was performed on ice in a stepwise approach where 1/10 fraction of streptavidin conjugate was added to the antigen followed by a ten-minute incubation. After the final incubation, 1 pmol free biotin was added and the mixture was incubated for 30 minutes on ice. Labelled antigens were stored at 4°C and used within two weeks.

To stain cells, PBMC were thawed with 50μg/mL DNAse I (Sigma) in pre-warmed RPMI + 10% FBS and washed once in media including DNAse I. Cells were then resuspended in PBS containing LIVE/DEAD® Fixable Violet Dead Cell Stain (ThermoFisher) with 10µg/mL purified streptavidin (to block aspecific binding, Biolegend) for 15 minutes. Then labelled antigens and an antibody cocktail containing CD71-AF700, CD19-BV605, CD27-PE/Cy7, CD38-APC/Cy7, CD69-BV510 and CCR10-APC (all Biolegend) was added and cells were incubated for another 15 minutes. Finally, cells were washed, resuspended in 200µL PBS and acquired on a LSR2.

### Immunohistochemistry

A nasal biopsy was fixed in 4% PFA for 16-24 hours before rinsing in 50% and 70% ethanol. This was embedded in Paraffin, cut into 4µm sections, dewaxed, subjected to antigen retrieval (95°C for 15 minutes in Sodium Citrate Buffer (pH 6) and processed for immunohistochemistry as published (Durrenberger et al., 2012). In short, sections were permeabilised in methanol for 15 minutes with 1% hydrogen peroxide. After rinsing in PBS, primary antibodies were diluted in goat (or horse) serum buffer (1% BSA, 4% goat (or horse) serum, 0.01% sodium azide in PBS). Primary antibodies used were: CD3 (Dako), CD4, CD20, CD66b, CD68, CD11b (Abcam), CD8 (Epitomics) and CD161 (Atlas antibodies), which were applied over night at 4°C. Sections were rinsed in PBS and secondary biotinylated antibodies (Vector lab) were applied for 45 mins at RT. Slides were rinsed and a complex of avidin and biotin (ABC) solution was added to sections for 60 minutes which was prepared 30 minutes prior incubation After rinsing, NovaRed™ (Vector®, Burlingame, CA, U.S.A) chromogen was prepared to manufacturer’s instructions. Sections were counterstained, dehydrated, placed in xylene and mounted for microscopy and scanned using the nanozoomer digital pathology (NDP, Hamamatsu, Photonics KK). Pictures were processed using the NDPview 2 software (version 2.6.13; Hamamatsu Photonics KK).

### ELISA

Serum IgG and IgA titres against *Streptococcus pneumoniae, Streptococcus pyogenes, Staphylococcus aureus* and *Haemophilus influenzae* were quantified in serum samples using whole cell ELISA. The ELISA was performed on MaxiSorp™ 96 well plates (Nunc). Per pathogen, 100μL of 10^8^ CFU/mL was prepared in carbonate buffer pH 8, added to the plates and allowed to adhere to the wells for 16 hours at 22°C. Then the plates were washed three times using phosphate buffered saline (PBS) containing 0.05% Tween 20, followed by blocking by adding 100 μL of PBS containing 2% Bovine serum albumin. Plates were incubated at 37°C for 1 hour and were washed before adding serial dilutions of serum samples. Standard curves for IgG and IgA were generated based on a standard pool serum (sera of 7 Spn carriers collected at D23 post challenge). Arbitrary units of IgG and IgA were assigned to the serum standard for each pathogen.

### Statistical analysis

Two-tailed, non-parametric statistical tests were used throughout the study. The number of cells in a cluster for each subject was normalized against the total number of non-immune cells acquired by CyTOF for that subject to account for number of cells isolated from a given biopsy. This normalization strategy has the advantage that the normalized frequencies of cells in a cluster is not dependent on other clusters, which is a major disadvantage of normalizing against total immune cells. Normalized cluster abundances were then compared between carriage^−^ and carriage^+^ subjects for each of the clusters using the Mann-Whitney test, without correcting for multiple testing. Data was analysed and graphs were created using ‘pheatmap’ and ‘ggplot2’ packages in R software and circular graph (Fig. 1C) was created using circos software (Krzywinski et al., 2009).

## Data availability

Normalized and debarcoded CyTOF fcs files have been deposited in the FlowRepository (https://flowrepository.org/).

## Supplemental information

**Supplementary Video 1.**
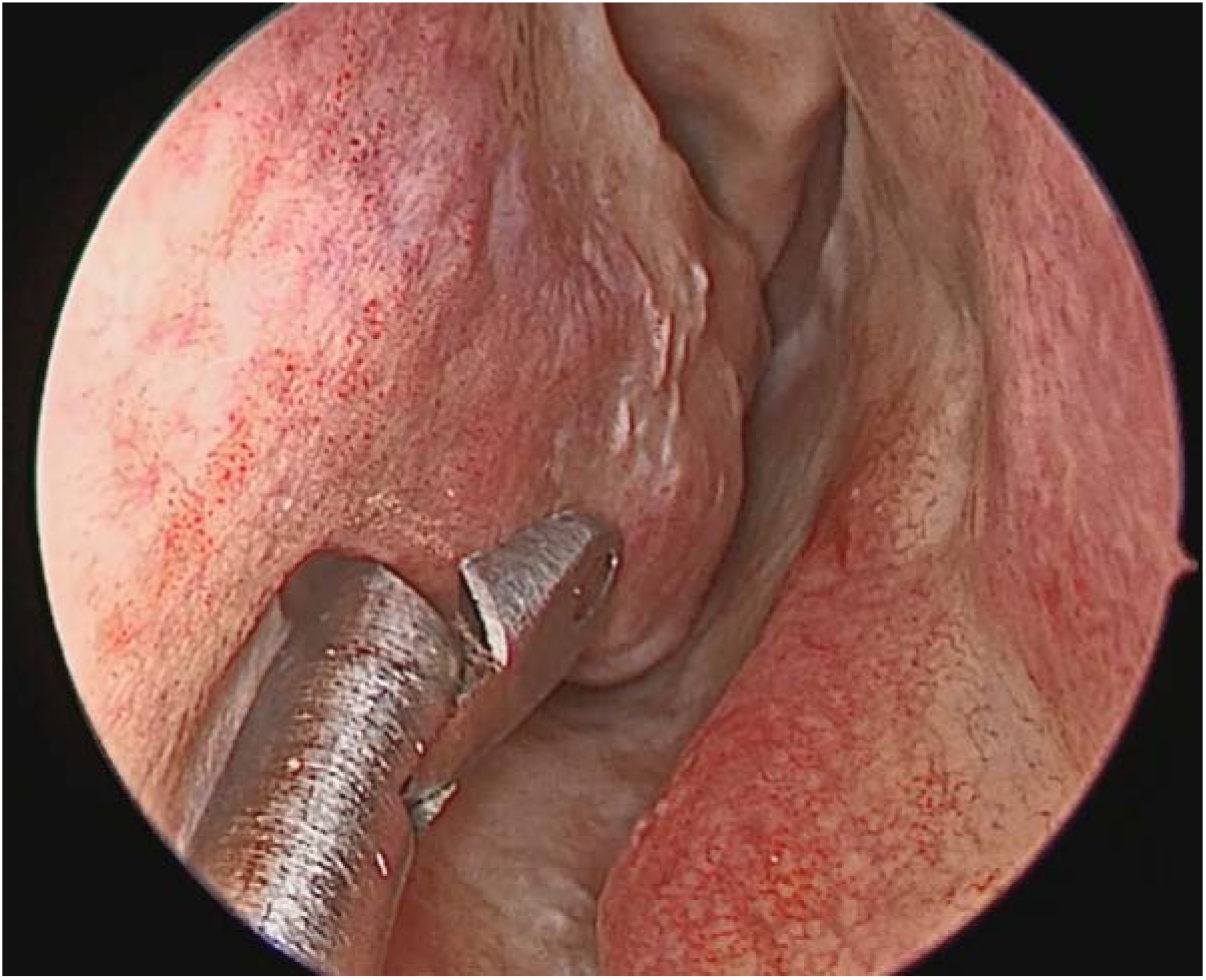
Video demonstrating the location and method of nasal biopsy collection.

**Supplementary Table 1.**
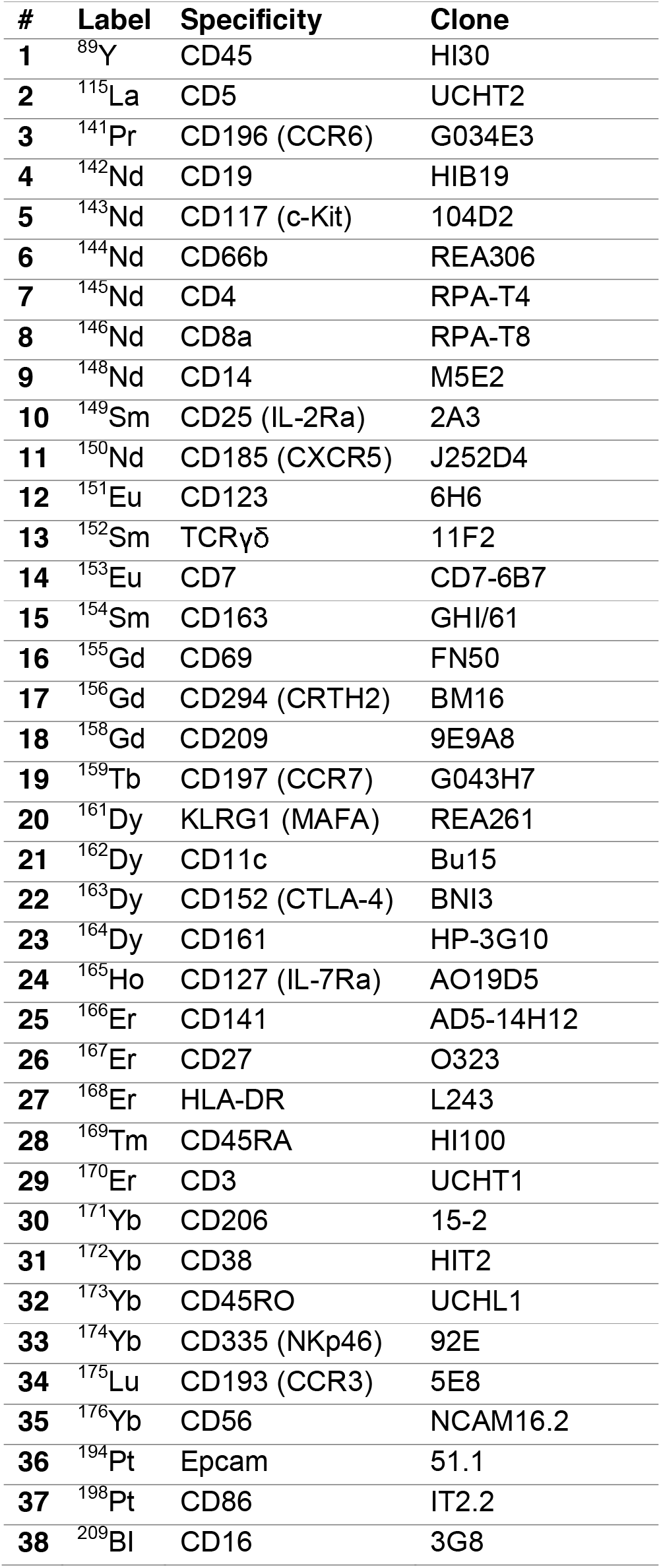
List of antibodies used for CyTOF.

| # | Label | Specificity | Clone |
| --- | --- | --- | --- |
| 1 | <sup>89</sup>Y | CD45 | HI30 |
| 2 | <sup>115</sup>La | CD5 | UCHT2 |
| 3 | <sup>141</sup>Pr | CD196 (CCR6) | G034E3 |
| 4 | <sup>142</sup>Nd | CD19 | HIB19 |
| 5 | <sup>143</sup>Nd | CD117 (c-Kit) | 104D2 |
| 6 | <sup>144</sup>Nd | CD66b | REA306 |
| 7 | <sup>145</sup>Nd | CD4 | RPA-T4 |
| 8 | <sup>146</sup>Nd | CD8a | RPA-T8 |
| 9 | <sup>148</sup>Nd | CD14 | M5E2 |
| 10 | <sup>149</sup>Sm | CD25 (IL-2Ra) | 2A3 |
| 11 | <sup>150</sup>Nd | CD185 (CXCR5) | J252D4 |
| 12 | <sup>151</sup>Eu | CD123 | 6H6 |
| 13 | <sup>152</sup>Sm | TCRγδ | 11F2 |
| 14 | <sup>153</sup>Eu | CD7 | CD7-6B7 |
| 15 | <sup>154</sup>Sm | CD163 | GHI/61 |
| 16 | <sup>155</sup>Gd | CD69 | FN50 |
| 17 | <sup>156</sup>Gd | CD294 (CRTH2) | BM16 |
| 18 | <sup>158</sup>Gd | CD209 | 9E9A8 |
| 19 | <sup>159</sup>Tb | CD197 (CCR7) | G043H7 |
| 20 | <sup>161</sup>Dy | KLRG1 (MAFA) | REA261 |
| 21 | <sup>162</sup>Dy | CD11c | Bu15 |
| 22 | <sup>163</sup>Dy | CD152 (CTLA-4) | BNI3 |
| 23 | <sup>164</sup>Dy | CD161 | HP-3G10 |
| 24 | <sup>165</sup>Ho | CD127 (IL-7Ra) | AO19D5 |
| 25 | <sup>166</sup>Er | CD141 | AD5-14H12 |
| 26 | <sup>167</sup>Er | CD27 | O323 |
| 27 | <sup>168</sup>Er | HLA-DR | L243 |
| 28 | <sup>169</sup>Tm | CD45RA | HI100 |
| 29 | <sup>170</sup>Er | CD3 | UCHT1 |
| 30 | <sup>171</sup>Yb | CD206 | 15-2 |
| 31 | <sup>172</sup>Yb | CD38 | HIT2 |
| 32 | <sup>173</sup>Yb | CD45RO | UCHL1 |
| 33 | <sup>174</sup>Yb | CD335 (NKp46) | 92E |
| 34 | <sup>175</sup>Lu | CD193 (CCR3) | 5E8 |
| 35 | <sup>176</sup>Yb | CD56 | NCAM16.2 |
| 36 | <sup>194</sup>Pt | Epcam | 51.1 |
| 37 | <sup>198</sup>Pt | CD86 | IT2.2 |
| 38 | <sup>209</sup>Bi | CD16 | 3G8 |

**Supplementary Figure 1.**
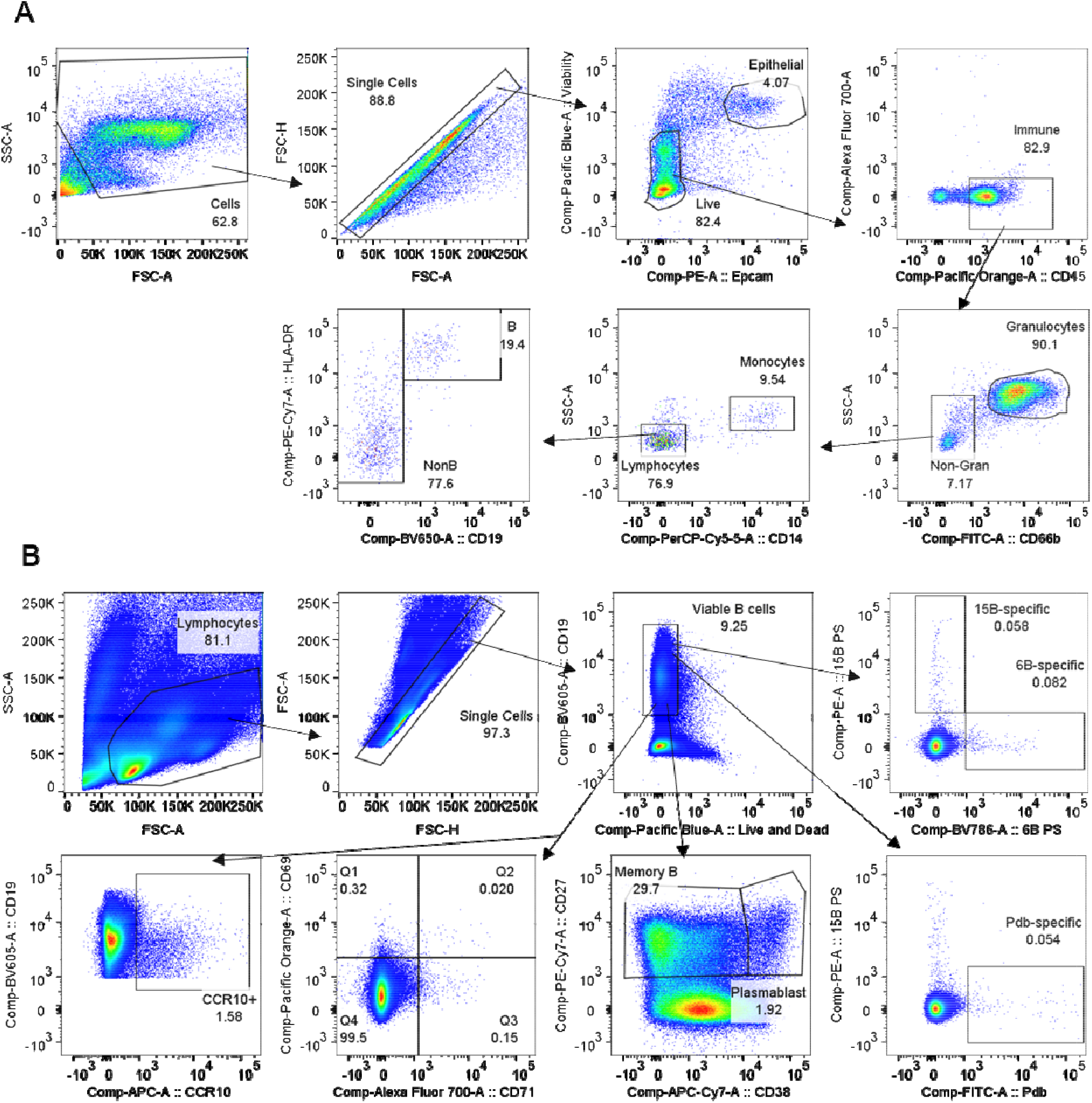
B cell validation flow cytometry gating strategies. A) Gating strategy for nasal CD19^+^ B cells for a representative nasal curette sample. Each plot shows the cells contained in the precedent gate. Population names and frequencies are shown. B) Gating strategy for the detection of pneumococcus-specific B cells. Each plot shows the cells contained in the precedent gate. Population and frequency are shown. Gates were set on all B cells and copied into Spn-specific populations.

**Supplementary Figure 2.**
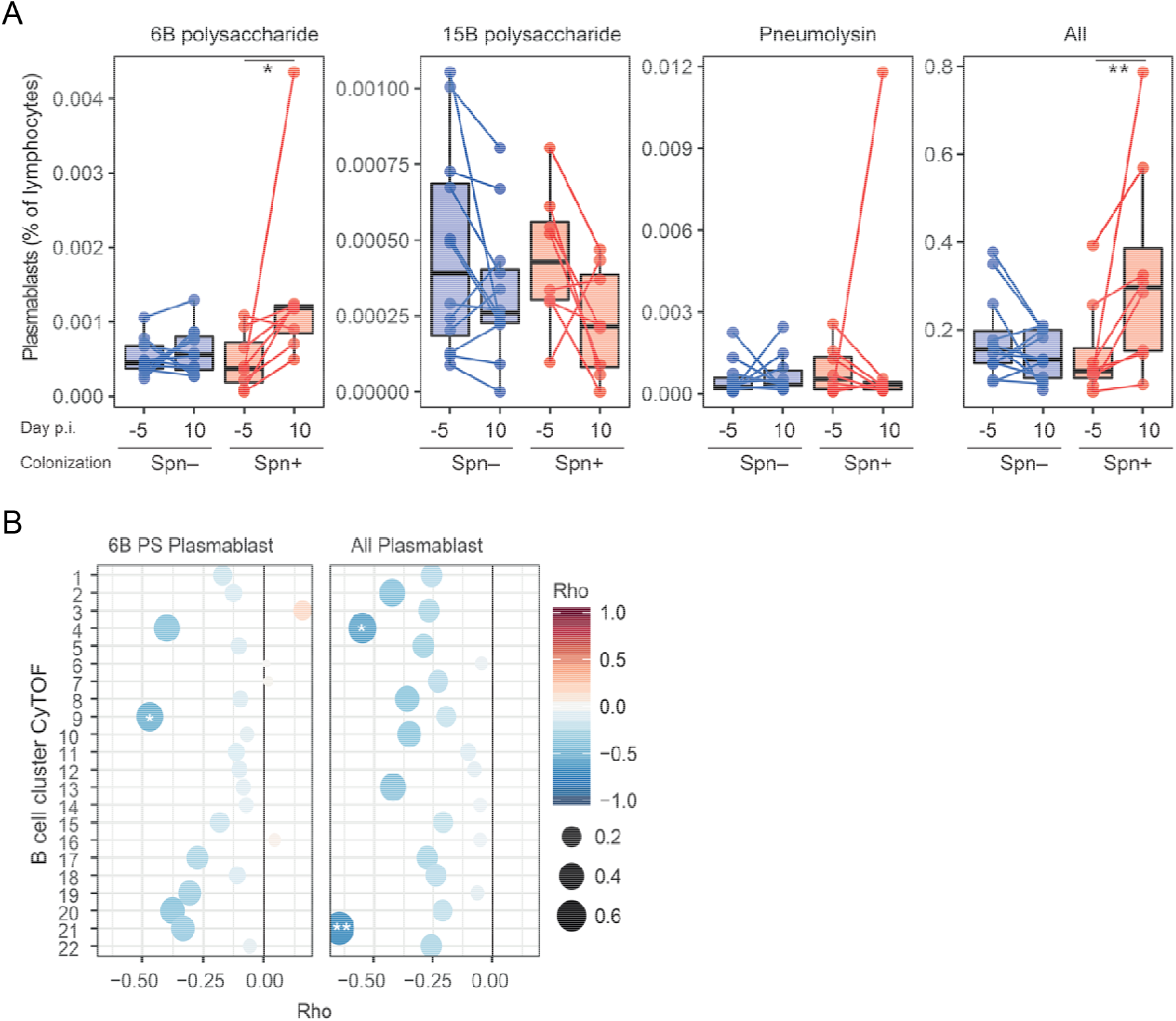
Normalization of circulating plasmablasts to total lymphocyte numbers. A) Levels of 6B polysaccharide-specific, 15B polysaccharide-specific, Pneumolysin derivative b (Pneumolysin)-specific or all plasmablast amongst total lymphocytes were measured from PBMC collected at baseline (Day -5) and at the time of biopsy (Day 10 post inoculation). Boxplots and individual subjects are depicted with carriage– in blue (n=12) and carriage+ in red (n=8), with paired samples connected by dashed lines. * p < 0.05, ** p < 0.01 by Wilcoxon test comparing a group to its baseline. B) Correlations between fold-change in levels of 6B PS-specific and total plasmablasts between baseline and day 10 normalized against total number of lymphocytes against levels of B cell clusters measured by CyTOF. Color and size of symbols reflect the Spearman rho value. * p < 0.05 and ** p < 0.01 by Spearman test.

**Supplementary Figure 3.**
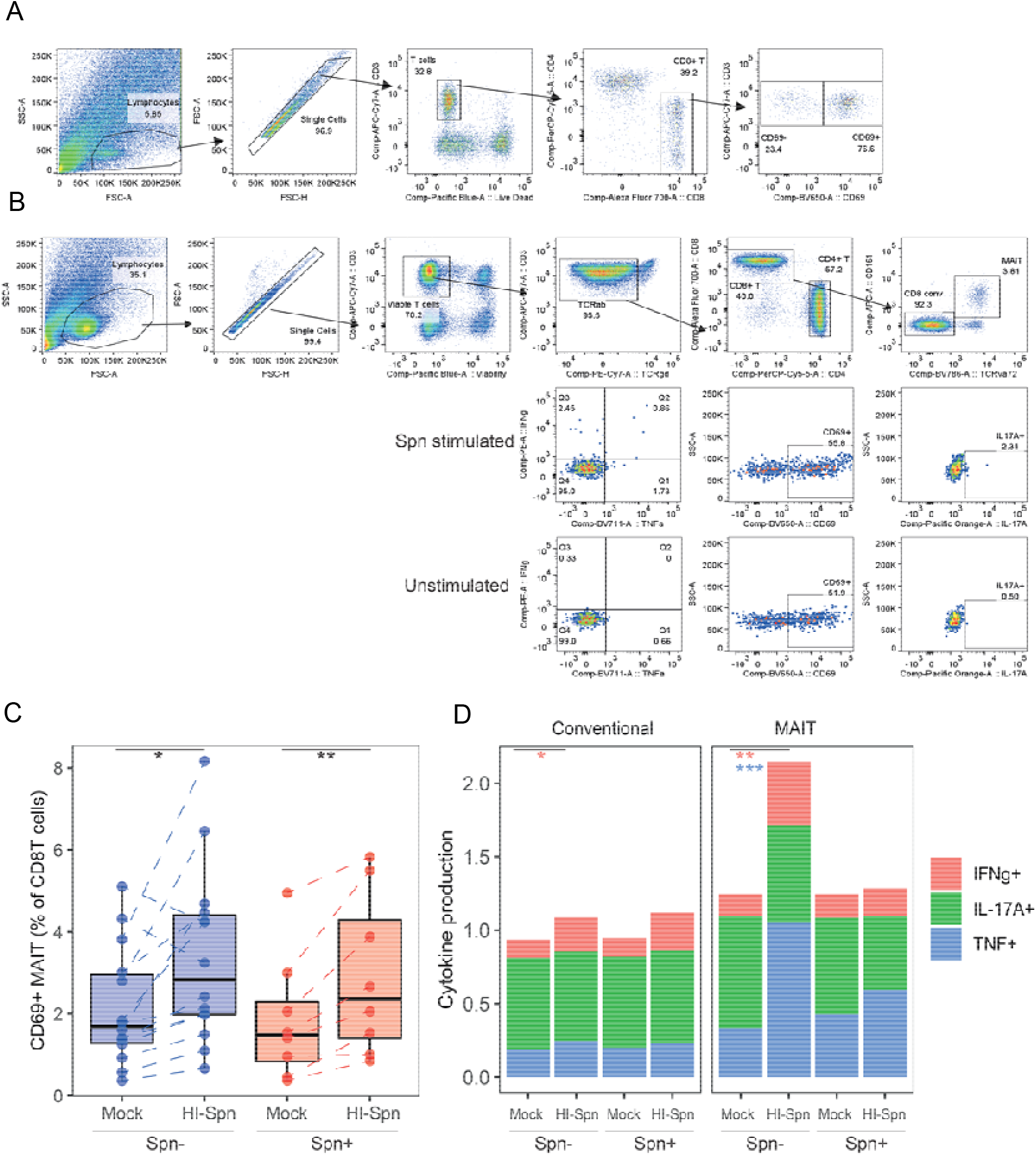
CD8^+^ T cell flow cytometry gating strategies. A) Gating strategy for CD8^+^ T tissue-resident memory cells by flow cytometry for a representative nasal biopsy. Each plot shows the cells contained in the precedent gate. Population and frequency are shown. B) Gating strategy to detect CD8^+^ mucosal associated invariant T cells in PBMC. One representative heat-inactivated pneumococcus (HI-Spn)-stimulated sample is depicted. For cytokine production and CD69 activation large dots are used to better show rare events. Each plot shows the cells contained in the precedent gate. Population and frequency are shown. C) Levels of CD69 on MAIT cells after stimulation or not. Individuals are shown and connected by lines and boxplots are overlaid. D) Stacked bar charts showing the median level of cytokine production (IFN-□ in red, IL-17A in green and TNF in blue) for conventional and MAIT CD8+ T cells. *p<0.05, **p<0.01, ***p<0.001 by Wilcoxon test.

**Supplementary Figure 4.**
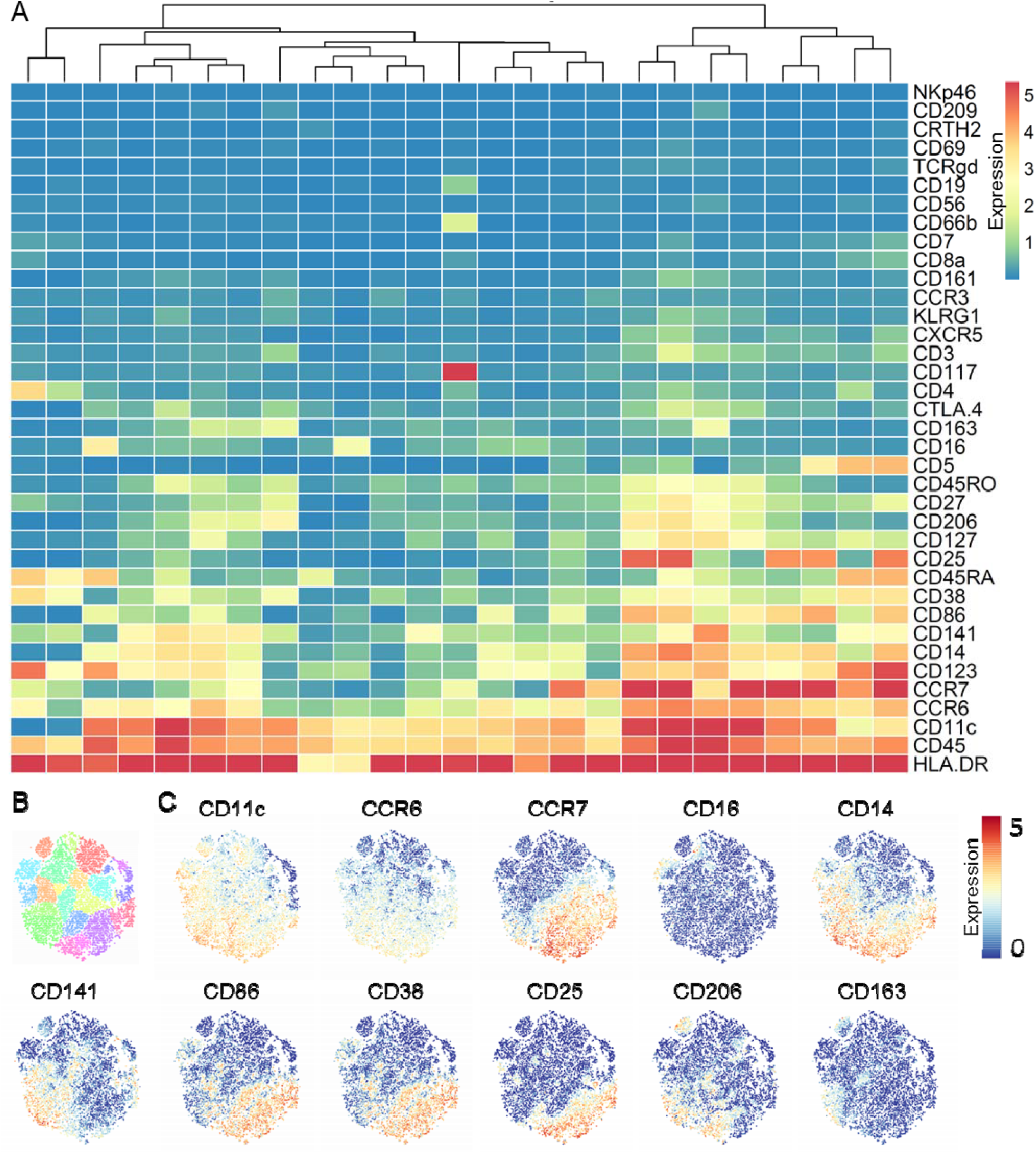
Nasal monocytes/macrophages are predominantly CD14^+^ CD16^−^ classical monocytes. A) Heatmap showing for each of the monocyte clusters (columns) the expression for thirty-seven markers (rows). Markers are ordered by increasing median expression. Columns are re-ordered and a cluster dendrogram is shown. B) Cluster definition within the myeloid cell lineage. C) Fingerprint graphs showing expression for selected markers on a single-cell level.

**Supplementary Figure 5.**
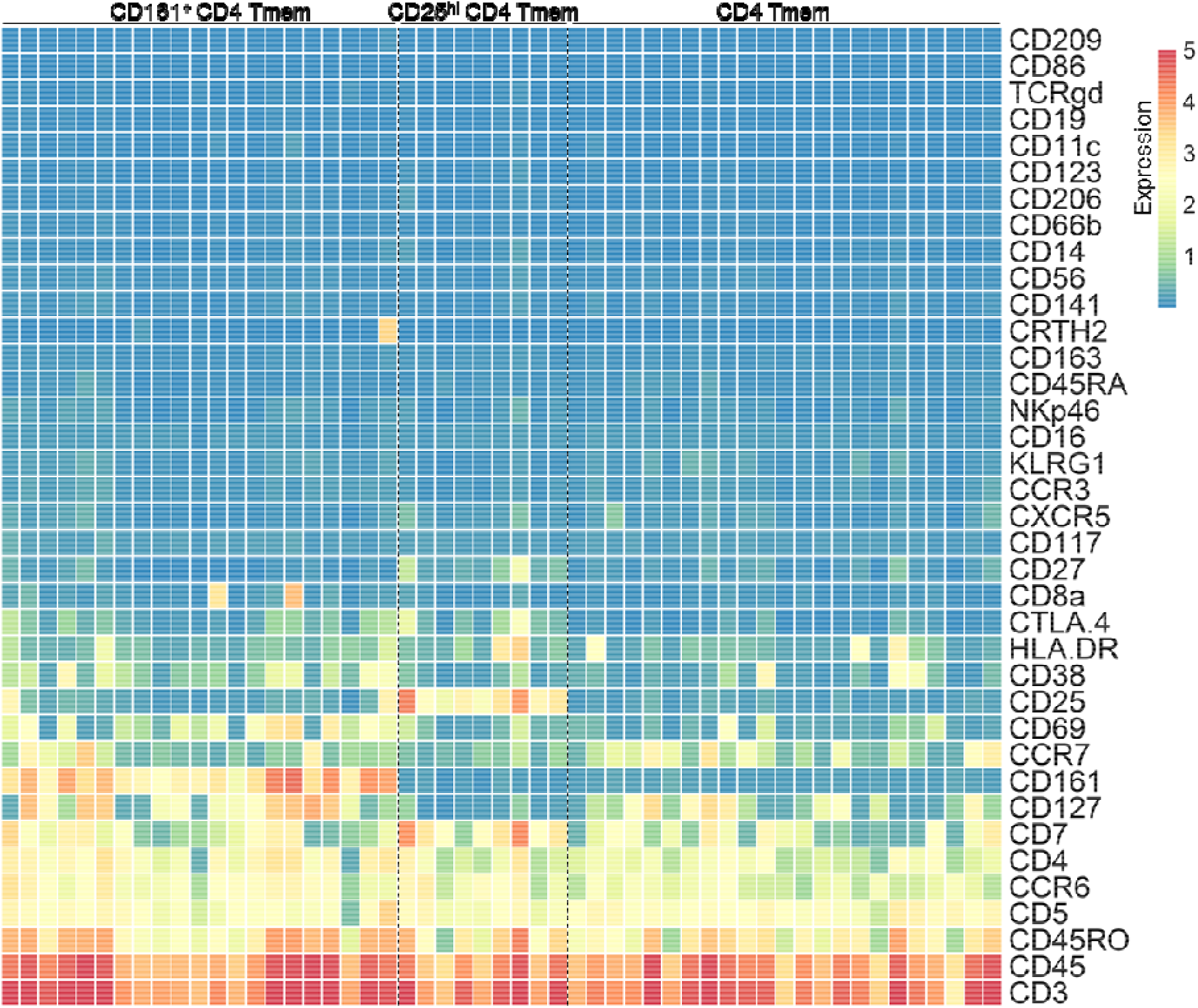
Nasal memory CD4 T cell phenotype. Heatmap including each of the three memory (CD45RO+) CD4+ T cell subpopulations within the CD4+ T cell lineage. All identified clusters (columns) with expression for thirty-seven markers (rows) are shown. Clusters belonging to the CD161^+^, CD25^hi^ and CD4 T memory (Tmem) subpopulations are separated by dashed lines. Markers are ordered by increasing median expression.

